# A General Method for Detection and Segmentation of Terrestrial Arthropods in Images

**DOI:** 10.1101/2025.04.08.647223

**Authors:** Asger Svenning, Guillaume Mougeot, Jamie Alison, Daphne Chevalier, Nisa Chavez Molina, Song-Quan Ong, Kim Bjerge, Juli Carrillo, Toke Thomas Høye, Quentin Geissmann

**Affiliations:** Department of Ecoscience, Faculty of Technical Sciences, Aarhus University; Center for Quantitative Genetics and Genomics, Faculty of Technical Sciences, Aarhus University; Department of Electrical and Computer Engineering, Faculty of Technical Sciences, Aarhus University; Centre for Sustainable Food Systems, Faculty of Land and Food Systems, University of British Columbia, Located on Traditional x^w^mƏθk^w^ƏýƏm Musqueam Territory

**Keywords:** Automated monitoring, insects, arthropods, entomology, deep learning, computer vision, multiple object detection and localization, instance segmentation

## Abstract

To better understand the status and trends of insects and other arthropods, emerging technologies like image recognition are developing rapidly. This is creating a strong demand for efficient and accurate algorithms for detection and localization of arthropods in images. Existing models have modest performance and do not generalise well to variation in scale, appearance and density of specimens, or imaging conditions. Consequently, each new application often requires manual labeling of training data and model training, which limits the uptake of image-based tools and technologies.

Here, we introduce `flatbug`, which is a powerful and general model to count and outline insects and other terrestrial arthropods in images. The training dataset is large and diverse and represent 23 different lab- and field-based imaging systems. The best `flatbug` model achieves an average *F* 1 = 94.2% on our validation dataset. Crucially, we show that `flatbug` has great out-of-the-box performance and generalises well to novel contexts. When images from a given dataset are left out of model training, the performance of `flatbug` is only reduced by on average 7.1% for the dataset in question.

By using truly stratified cross-validation, we set a precedent for robust evaluation of deep learning model performance and generalization. We also take steps towards scale- and size-agnostic arthropod detection, by developing an integrated tiling framework for inference and training. Additionally, `flatbug` ‘s implementation of `YOLOv8` for instance segmentation enables downstream background removal and body size estimation.

The generaliseability of `flatbug` stems from the diversity of contexts represented in the `flatbug` dataset, including 113550 arthropods annotated across 6131 images. Alongside a fully documented Python package with tutorials for integration and analysis via https://github.com/darsa-group/flat-bug/, the `flatbug` dataset is available from https://www.doi.org/10.5281/zenodo.14761447. By providing performant models and the accompanying dataset, `flatbug` offers both a ready-to-use tool and a benchmark for the future. Overall, `flatbug` represents a significant methodological advance within arthropod image detection, with user-friendly integration for monitoring and research.

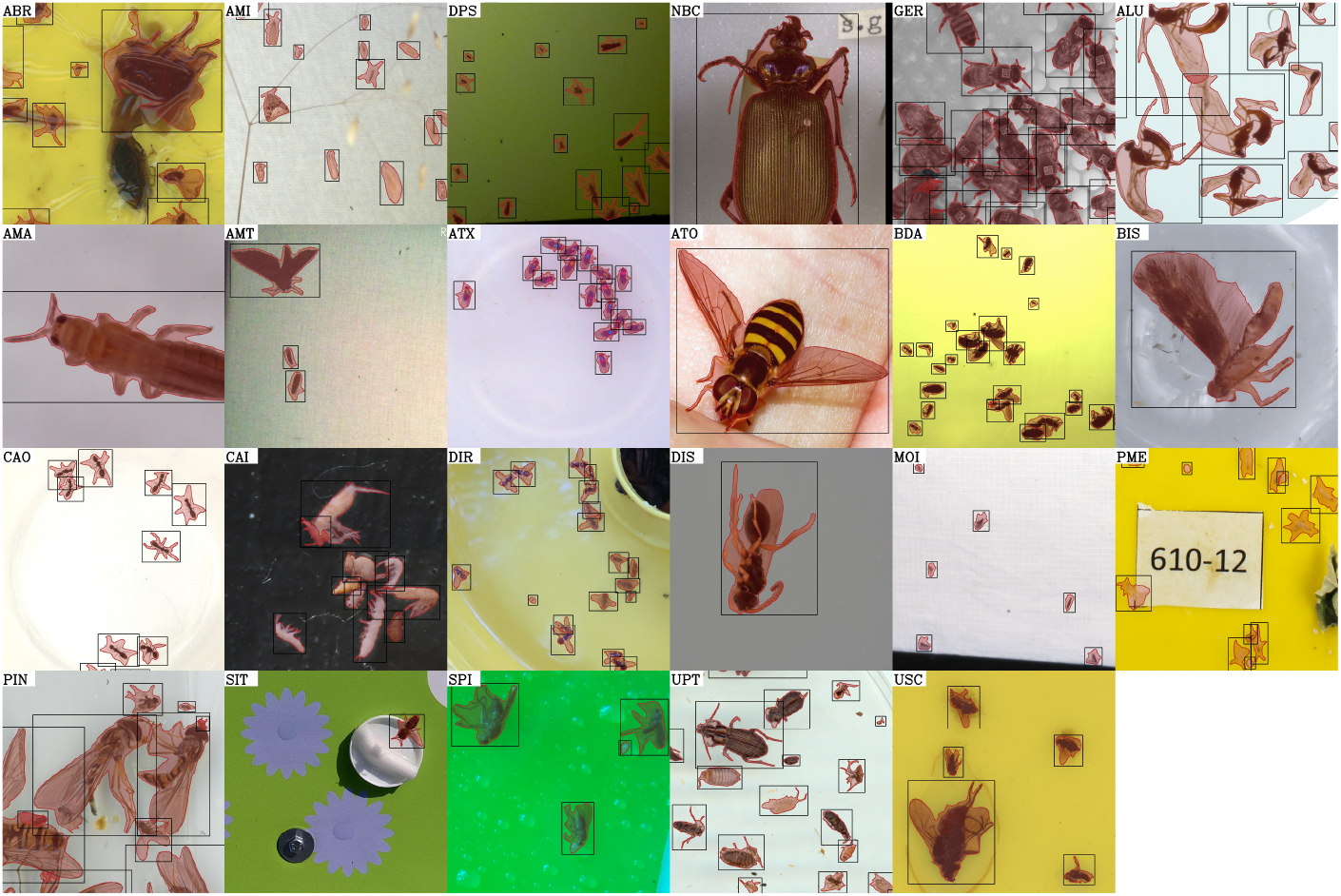

## 1 Introduction

Arthropods are an ecologically important group of organisms under pressure from multiple global change drivers (Wagner et al., 2021). Therefore, it is critically important to understand which arthropods are affected the most and by which drivers, where and when. However, for most arthropod taxa this is currently not possible due to the acute need for better and more standardised arthropod monitoring data, which consequently delays the production of ecological knowledge and conservation actions (Thomas et al., 2019). An important constraint is that traditional arthropod monitoring relies heavily on manual counting and identification of specimens, which requires highly specialised expertise, is very time-consuming and thus costly.

Recently, novel image-based tools for entomology have begun to emerge (Alison & Høye, 2024; Gal, Saragosti, & Kronauer, 2020; Geissmann, 2022; Gharaee, Gong, Pellegrino, Zarubiieva, Haurum, Lowe, McKeown, et al., 2023; Høye et al., 2021; van Klink et al., 2022). However, most of these tools depend on accurate localisation of individual insects in images, and it has been recognised that such tools need to be improved (Høye et al., 2025; van Klink et al., 2024). Therefore, general, powerful and accurate machine learning-based tools are needed for automatically detecting and extracting individual arthropods on a broad spectrum of image types, backgrounds, and resolutions. This task is particularly difficult due to the diversity and complexity of possible images in arthropod monitoring contexts, encompassing multiple orders of magnitude in both image and organism size, image-to-organism size ratio, and organism density, as well as variable lighting, imaging conditions and background compositions.

Generalized machine learning models have already begun to transform monitoring of larger fauna and flora, with tools such as MegaDetector (Beery et al., n.d.) and iNaturalist (“A Community for Naturalists · iNaturalist”, n.d.), among others, enhancing scalability and reducing costs for species detection and identification across mammals (Aodha et al., 2018; Shepley et al., 2021; Willi et al., 2019; Z. Wu et al., 2023), amphibians (Kimura & Sota, 2023), birds (Stowell et al., 2019), insects (Hong et al., 2021), plants (Mäder et al., 2021; “Pl@ntNet”, n.d.) and more (Kirillov et al., 2023; Kloster et al., 2023). However, existing methods for arthropod detection are typically developed for specialized use cases and lack the ability to generalize across diverse taxa and contexts (Hong et al., 2021; Schneider et al., 2023; Sys et al., 2022). Some previous works, such as Mazen (2023), have used citizen data to create a general model; however, despite the diversity of citizen science images, they rarely contain more than one individual and are not standardized. The lack of open and standardized datasets adhering to the FAIR (Wilkinson et al., 2016) principles (Findable, Accessible, Interoperable, Reusable) has slowed progress in data annotation and decreased its efficient usage (Schneider et al., 2023; van Klink et al., 2022).

Although many models, such as the YOLO (Redmon et al., 2016) series and Mask R-CNN (He et al., 2018), are widely applied for generalized multiple object detection (and segmentation), these models are typically limited to fixed-size inference at relatively low resolutions, making them unsuitable for processing large images containing small instances. To overcome this challenge, the hyper-inference framework SAHI (Akyon et al., 2022) is often applied on top of models such as YOLOv8 (Jocher et al., 2023). SAHI performs hyper-inference— inference with a base model using a meta strategy—by tiling the image in fixed size tiles at the native scale, with some specified overlap, and then combines the tile predictions with a full-image prediction. This enables the usage of fixed-size inference models such as the YOLO series and Mask R-CNN for large images with instances at the native and full scale, but falls short for intermediate-size instances, especially for very large images.

In addition to addressing fixed-size inference on large images, many detection models also contend with duplicate predictions, a problem which is only exacerbated by SAHI-like hyper-inference. In practice, SAHI, the YOLO series(Redmon et al., 2016), and others tackle this challenge by applying prediction deduplication (or merging) through non-maximum suppression (or its variants), which computes the Intersection over Union (IoU) on bounding boxes rather than on instance segmentation polygons or masks—a strategy frequently used even in segmentation models because calculating intersections for concave polygons is considerably more computationally expensive than for axis-aligned rectangles.

Most current arthropod detection models rely on bounding boxes—either exclusively for both training and inference or at least for processes like non-maximum suppression—primarily due to the lower annotation cost and reduced computational demands compared to segmentation. However, while instance segmentation requires more labor-intensive annotations and involves more complex IoU computations (e.g., between polygons), it offers several crucial benefits: it facilitates more accurate deduplication by leveraging precise object boundaries, enables effective background removal, and improves instance size estimation. These advantages are key to advancing automated arthropod monitoring.

### 1.1 Contribution

In this manuscript, we aim to address the above challenges in arthropod detection, and provide a practical tool for monitoring schemes, lab experiments, and related applications. Additionally, we aim to set a precedent for more reproducible, effective, robust, and general models and metrics within the machine learning and arthropod research communities, with emphasis on data sharing and evaluating model generalization.

To that end, we developed `flatbug` (https://github.com/darsa-group/flat-bug/), a general multiple-object detection and segmentation model for terrestrial arthropods recorded by diverse imaging systems. `flatbug` is an open-source model, but also a unified framework for training, evaluation, and inference, built on YOLOv8 (Jocher et al., 2023) and trained on a combination of entirely novel data and re-annotated datasets from the literature (Geissmann & Svenning, 2025). `flatbug` integrates a custom-built, scale-agnostic hyper-inference algorithm inspired by SAHI (Akyon et al., 2022), with an appropriate training domain and an evaluation pipeline, specifically tailored for training YOLOv8 models intended for use with `flatbug`. Crucially, in contrast to YOLOv8 and SAHI, `flatbug` principally employs a segmentation-first approach: using segmentation polygon IoU (Intersection over Union) for deduplication, which compares the overlap between the actual shapes of detected objects, rather than their bounding boxes, and is therefore better able to accurately distinguish closely spaced individuals, enhancing `flatbug`’s ability to detect tightly packed arthropods.

As well as fully accessible, open code for our model and hyper-inference algorithm, we provide the accompanying training and validation dataset following the FAIR principles (Wilkinson et al., 2016). The dataset for `flatbug` is unprecedented in its diversity, utilizing 23 subdatasets from distinct arthropod detection applications. Crucially, the dataset is explicitly stratified, allowing our approach to reach unprecedented performance by deploying a true novel-data evaluation regime. Rarely deployed in previous work, this mitigates data leakage and provides more accurate assessments of generalization. Exploiting the diversity and explicit stratification of the `flatbug` dataset, we take a step beyond previous efforts to perform a set of robust cross-validation experiments. First, we explore how removing a subdataset from training affects performance for all datasets in a typical leave-one-out cross-validation procedure. Furthermore, we consider second-order interactions between individual subdatasets (strata) using an original leave-two-out cross-validation approach, allowing us to investigate the relationships and redundancy between subdatasets using a novel inter-strata redundancy metric.

## 2 Methods & Materials

All experiments, models and software are built in `Python 3.11`, while experiment statistics and figures are created in `R 4.4.1` using the `tidyverse` extensively (R Core Team, 2024; Van Rossum & Drake, 2009; Wickham et al., 2019). Deep learning modeling and post-processing is based on `PyTorch`, `NumPy` and the YOLOv8 segmentation model architecture, while some polygon calculations are executed with `shapely` (Gillies et al., 2025; Harris et al., 2020; Jocher et al., 2023; Paszke et al., 2017). Manual annotations were performed using the online annotation platform `CVAT` (Sekachev et al., 2020). All training experiments were run on Ucloud through DeIC on NVIDIA H100 hardware in Linux Ubuntu 24.04 using `submitit` and SLURM for process management(“Facebookincubator/Submitit”, 2025; Yoo et al., 2003).

### 2.1 Dataset

We constructed a diverse dataset by compiling a set of 23 subdatasets (either publicly available images or constructed for this article) covering a wide domain in terms of arthropod taxonomy, cameras, pose and background. In most cases, since most original images were not annotated for instance segmentation, we annotated these images manually, or semi-automatically. In addition to re-annotating available images for 15 subdatasets derived from existing data, we produced 8 previously undescribed subdatasets. The resulting dataset, containing 6131 annotated images from the 23 subdatasets, is available under the Creative Commons license (10.5281/zenodo.14761447) (Geissmann & Svenning, 2025). Table 1, lists, references and briefly describes the individual subdatasets. Each subdataset has its own DOI and is deposited as a separate Zenodo entry which can be found in Table S1.

**Table 1:**
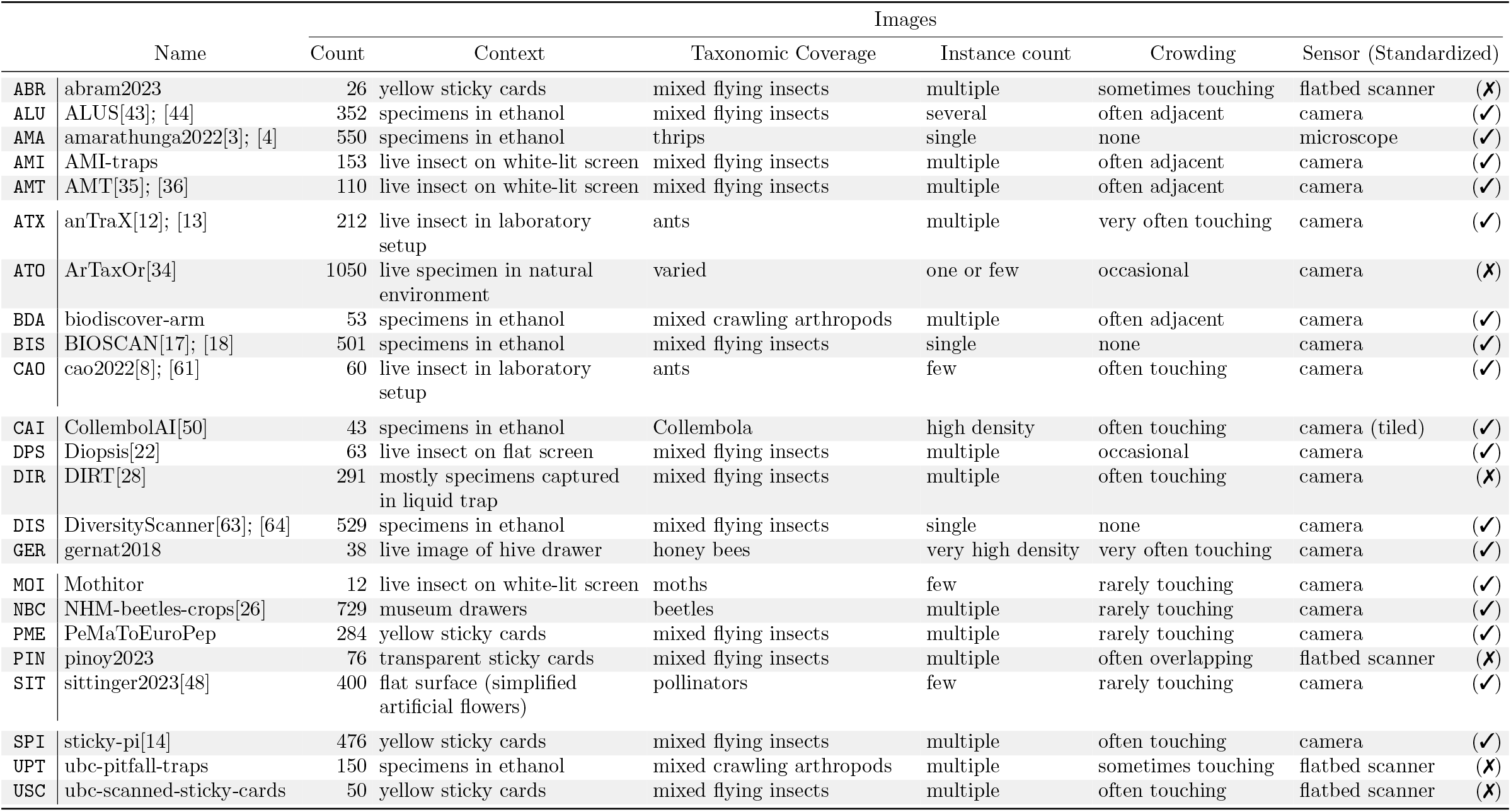
Dataset description and characteristics. The three-letter abbreviation and full names of each subdataset can be found in the first two columns, while image charactistics and content descriptions can be found in the remaining six columns.

### 2.2 Training

`flatbug` training employs a modified YOLOv8 segmentation training regime, with changes restricted to the sampling and image augmentation stages. Training images are sampled in a manner that mirrors the hyper-inference prediction regime, which the models are intended for. This is achieved by sampling ‘tile’-like crops from the full images in the dataset using the following strategy: First, we sample a relevant zoom level, *Z*:

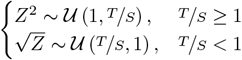

where *T* is the desired tile size (default=1024) and *S* = max{|*D*_*x*_|, |*D*_*y*_|} is the maximum dimension size (height; width) of the image. The square root or power of two is added to adjust for the number of possible unique tiles at each zoom-level. Given a specific zoom level *z* realized from *Z*, we then either sample or calculate a coordinate offset for the tile *o* (if the tile size is larger than the dimension, we center the tile):

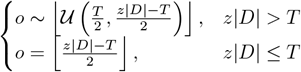

for each dimension. After extracting a random tile from an image, we perform the following standard augmentation operations; rotation (uniform degree probability), color jitter, color inversion (25% probability) and horizontal/vertical flip. The color jitter, horizontal, and vertical flip parameters are controlled through the standard YOLOv8 hyperparameter interface, while the translate and scale parameters from YOLOv8 have been disabled (since they conflict with our tile sampling strategy). We then remove instances with less than 97.5% or 32px of their (scaled) area within the extracted image region (tile)—either because they are fully/partially outside the region, or because they are too small at the sampled zoom level. The removed instances are inpainted using the OpenCV(Bradski, 2000) ‘telea’ inpainting method(Telea, 2004) if they overlap with the image region to avoid obvious inpainting artifacts. Secondly, we deterministically oversample images to account for the above process: larger images have many more potentially unique ‘tiles’ approximately proportional to the pixel area of the image. This step ensures that the training process is not biased by differences in how each subdataset is stored; some subdatasets consist of ‘precomputed’ tiles, where the original images have been gridded and cut before annotation, leading to these subdatasets containing many more individual images than other subdatasets where the full images are annotated instead. See Appendix K for the technical details.

### 2.3 Inference

`flatbug` employs a custom-built hyper-inference pyramid-tiling algorithm—visualized in Fig. 1 and Fig. 2— comprising pre-processing, tiling, and inference with post-processing. Before inference, the image is zero-padded by 32px on all sides to avoid edge artifacts and ensure consistent inference coverage, particularly near boundaries. Tiling begins by defining *N*_*S*_ zoom levels, *Z*, as a geometric series running from

**Figure 1:**
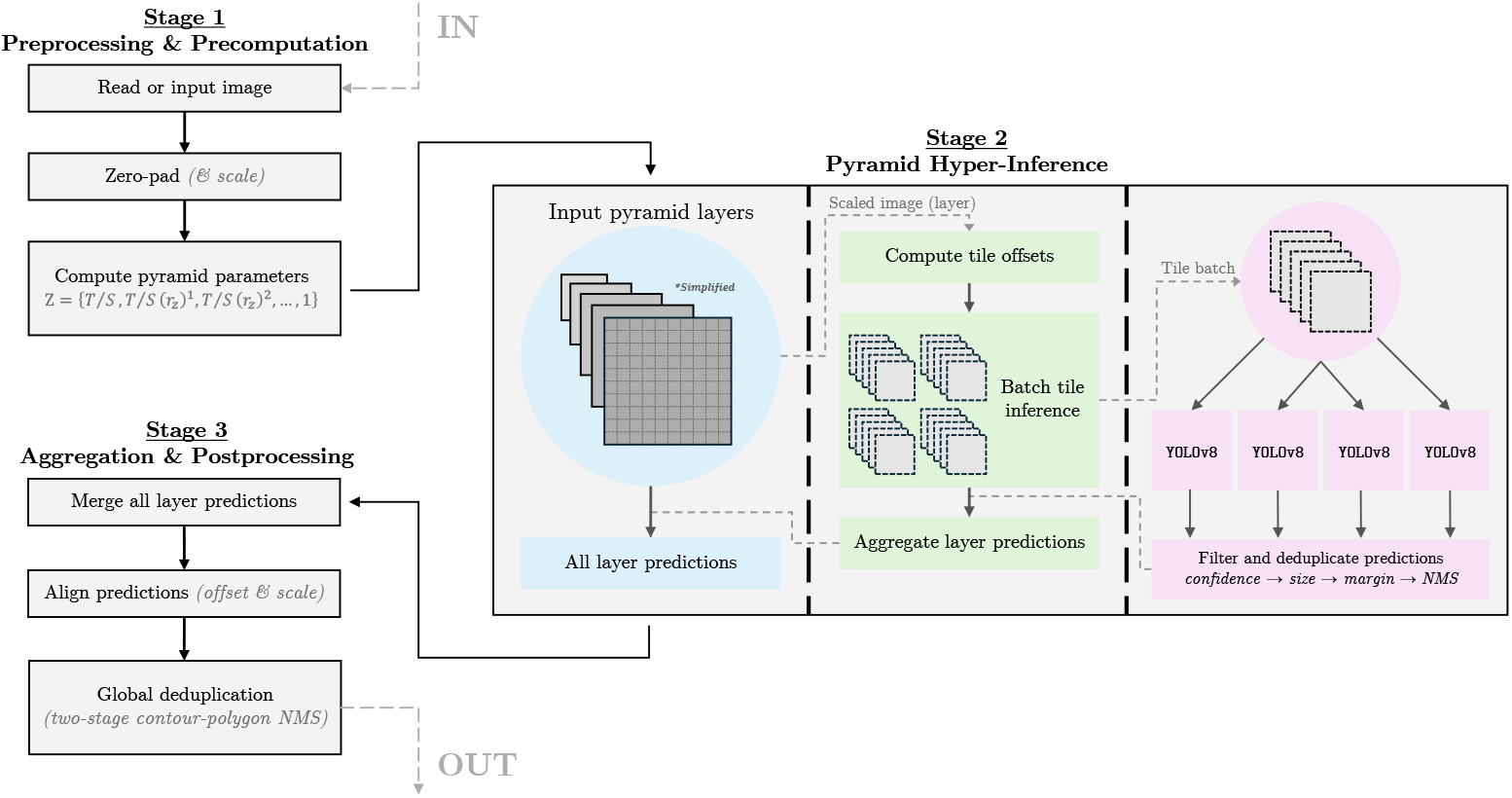
Simplified visual diagram of the `flatbug` inference pipeline. The `flatbug` inference pipeline is fully GPU-accelerated for all steps between reading the input image and global deduplication with CUDA and half-precision compatibility. Global deduplication is also mostly GPU-accelerated, however it proved simpler to use `shapely`(Gillies et al., 2025) a Python wrapper around `GEOS`(GEOS contributors, 2024) for polygon intersection calculation specifically. Only steps we deemed semantically or technically important in highlighting the defining characteristics of the pipeline are included. The code is available on the `flatbug` repository (https://github.com/darsa-group/flat-bug/). See Fig. 2 for an accurate depiction of the tile pyramid.

**Figure 2:**
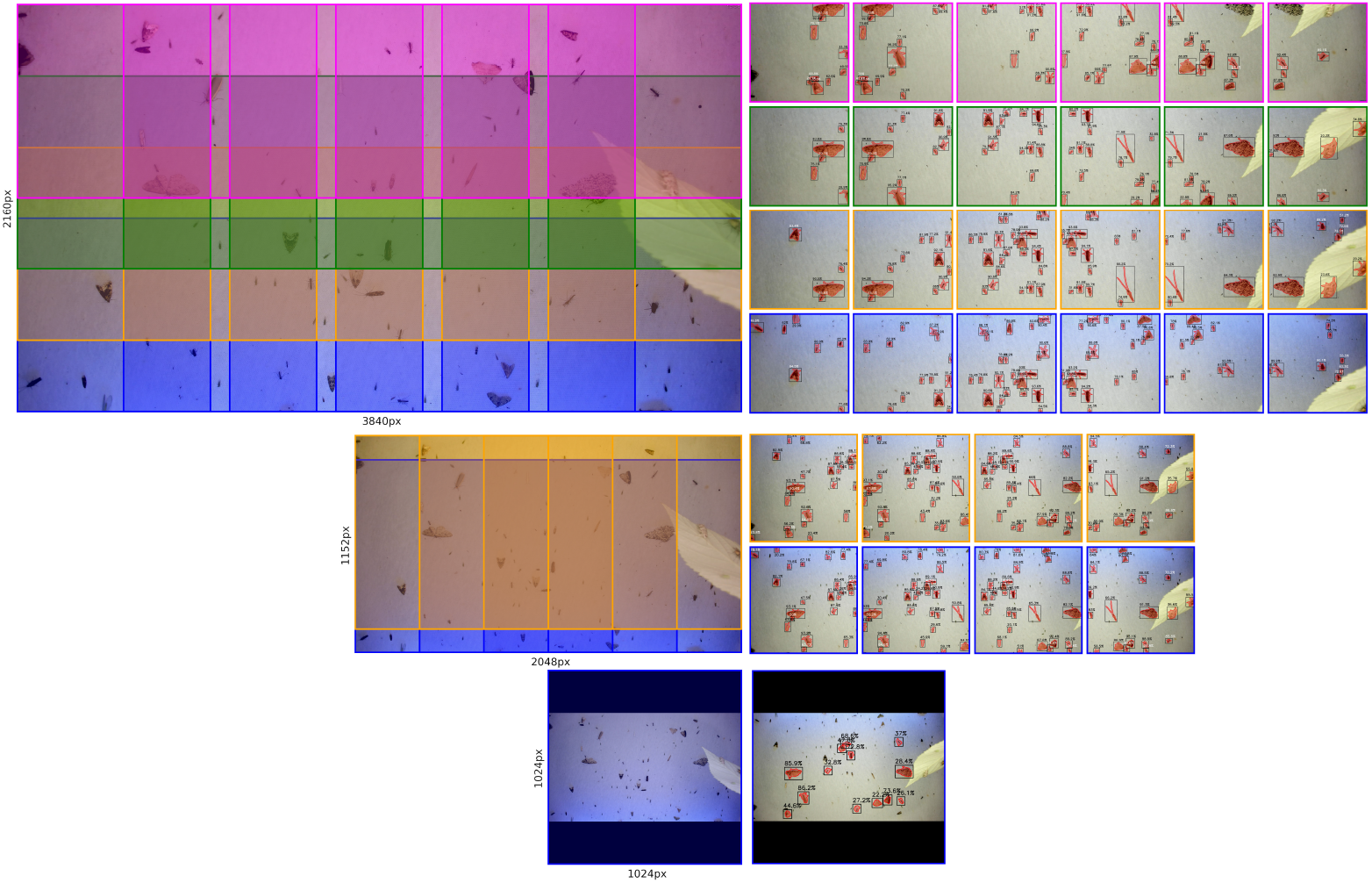
Example visualization of our pyramid-tiling strategy on an image from the AMI subdataset. Each row corresponds to a ‘layer’ in the pyramid. Left: Tiles overlayed on the scaled image with scaled dimension sizes on the left/bottom. Right: Extracted tiles with predictions; all tiles are 1024px × 1024px.

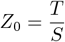

to

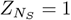

with a common ratio of *r*_*Z*_ = 1.5 by default. For efficiency we truncate the zoom levels at 0.9 and always include 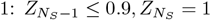. For each zoom level, a minimal list of tiles is generated based on a specified tile size and minimum overlap, ensuring overlaps are almost equal (± 1 pixel). The top (*y*) or left (*x*) corner coordinate of each tile, *p*, is calculated independently for each dimension using:

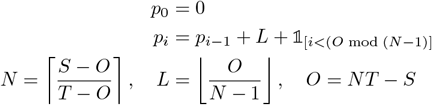

where *L* is the step size, *O* represents the total overlap, and *N* is the number of tiles required to cover a dimension of size *S* with tiles of size *T*. This formula ensures that non-divisible overlaps are allocated 1 pixel at a time to initial tiles, creating a consistent layout. The tile top-left corner coordinates, *P*, are then given by all pairs of corner top and left coordinates for each dimension:

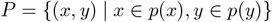

If the scaled and padded image is smaller than the tile size in a dimension, *T < S*, then this dimension is zero-padded symmetrically such that a single tile exactly covers this dimension, *T* = *S*.

Then, given a zoom level, *Z*_*i*_, and the corresponding tile positions, *P*, tiles are processed in batches (default=16) using YOLOv8 inference with two key post-processing modifications: First, bounding box IoU is replaced with contour polygon IoU for deduplication, improving performance on irregular shapes. Second, predictions near tile edges (default: ≤16px) are removed to reduce detections of instances that are artifically clipped. The predictions are then offset and scaled to align with the tile positions and zoom levels before combining the results across all zoom levels. Finally, positions are adjusted to account for the initial image padding, and global deduplication is performed using a two-stage NMS (non-max suppression) approach similar to Tan and Wang (2024). The first stage reduces the number of polygon intersection calculations by identifying connected components via pairwise bounding box IoU with a reduced threshold (default=0.05). The second stage applies standard polygon-based NMS within each component, reducing the number of polygon intersection calculations from *O*(*N* ^2^), where *N* is the number of predicted instances, to

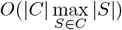

where *C* is the set of connected components. This approach drastically improves efficiency in images with low instance density, since it is significantly faster to calculate intersections between axis-aligned rectangles (bounding boxes) than arbitrary (simple) 2D polygons.

### 2.4 Evaluation

For comprehensive model evaluation, we used a simple custom-built end-to-end evaluation scheme on the fixed validation split. The end-to-end evaluation starts by running full hyper-inference on all images in the validation split. For each image, we then attempt to match ground-truth instances with predicted instances using a modified NMS algorithm (for tie-breaking) and a polygon-IoU threshold (default=0.2). Then we calculate bootstrap summary statistics recall, precision, and F1 with percentile confidence intervals, denoted as *Q*_*x*%_(*F*^*T*^) where *x*% is the percentile, *F* is the statistic, and *T* is the treatment or stratum, for each subdataset included in the stratum given by *T*. The bootstrapping procedure simply resamples instances—either inferred/FP (false positives) or labels/FN (false negatives) or matched/TP (true positives)—stratified by subdataset. Changes in performance of a metric are either denoted by 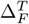 for the absolute change in statistic *F* or 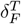 which gives the normalized relative change in statistic F, on only the images in the stratum given by *T* :

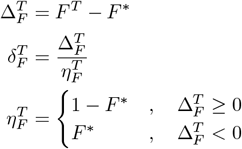

where *F* ^*^ is the baseline (control) under the specified treatment and/or stratum, *T*. The normalization factor *η* for *δ*simply ensures that *δ*∈ [−1, 1] in contrary to the absolute change with the range Δ ∈ [−*F* ^*^, 1 − *F* ^*^]. Intuitively, this means that if *δ*= −1*/*2 (−50%) then the treated model has gained 50% of the potential performance gain, and if *δ*= −1*/*2 (−50%) then it has lost 50% of the potential performance loss.

For all evaluation metrics we also make sure to remove all ground-truth and predictions with below 32px, as measured by the square root of the pixel are of the contour polygon, as these would also have been removed during training. See Appendix N for further details.

For in-training diagnostics, we utilize the native YOLOv8 segmentation evaluation scheme with the previously described modified training augmentation and oversampling techniques on a fixed validation set five times, since our modified augmentation (potentially) samples a ‘tile’ which covers a smaller region of each image. Validation is conducted after the first epoch, then every 5 epochs by default, and then once at the end of training.

### 2.5 Experiments

All experiments comprised two sequential steps: (1) training and (2) end-to-end evaluation. The following training hyperparameters are shared for all experiments: initial learning rate (lr0 = 10^−3^), final learning rate proportion (lrf = 10^−5^), stochastic gradient descent (optim = SGD), and batch size (batch = 32), as well as all other YOLOv8 segmentation training hyperparameters not mentioned here. The three ‘experiments’ were as follows: (1) training YOLOv8 segmentation models at the four available sizes nano (N), small (S), medium (M) and large (L), (2) leave-one-out cross-validation training with fine tuning, and (3) leave-two-out cross-validation training. These experiments were designed to quantify and evaluate the performance of `flatbug` using the four available YOLOv8 backbone sizes, the generalizability of `flatbug` to new images both similar and dissimilar to the images in the subdatasets of the `flatbug` dataset, and the relationships between the subdatasets across the domain of possible subdatasets. Experiments 2 and 3 were performed using only the medium (M) YOLOv8 segmentation model variant for 50 and 20 epochs, respectively, while models in experiment 1 were trained for 300 epochs. For efficiency, we only performed in-training evaluation every 25 epochs during our experiments. Further details for each experiment are presented below.

#### 2.5.1 Experiment 1: Backbone size comparison

Four models, one at each of the available model sizes in YOLOv8 (L/M/S/N), were trained for 300 epochs on our entire training dataset and evaluated on the entire validation dataset.

#### 2.5.2 Experiment 2: Leave-one-out cross-validation

24 medium (M) size models were trained for 50 epochs, 23 out of 24 models (corresponding to the 23 subdatasets) were trained on our training dataset with a particular subdataset left out, while the last model was trained on the full dataset as a baseline (control); these are the OOB (out-of-box) models. Each of these models was then trained for a further 10 epochs only on the subdataset that was left out (the control model was again trained on the full dataset); these are the FT (fine-tuned) models. For this experiment (Section 3.2) we only present changes in performance metrics at the level of individual subdatasets. In other words, we assess how excluding a given subdataset from training affects the end-to-end evaluation metrics for that same subdataset. The baseline (control) metrics were taken from the two full dataset models (OOB and FT) evaluated on each subdataset.

#### 2.5.3 Experiment 3: Leave-two-out cross-validation

A subset of 16 out of the 23 subdatasets were chosen (ALU, AMA, AMI, AMT, ATO, BDA, BIS, CAI, DIS, DPS, GER, PIN, SIT, SPI, UPT, USC) for leave-two-out cross-validation. The subset of 16 subdatasets were chosen ad-hoc, aiming to be as representative of the full 23 subdatasets as possible, while reducing the number of sub-experiments (i.e. models) to a manageable number. For each unique pairwise combination (including pairs of the same dataset) we trained a medium (M) model for 20 epochs on all 23 subdatasets except the particular pair of subdatasets. A control model was also trained for 20 epochs on all 23 training datasets. Subsequently, all resulting models (N=136 + 1) were evaluated on all datasets leading to a data-cube for each validation metric, **F**, where element **F**_(*k*)*i,j*_ corresponds to the value for metric *F* only on subdataset *k* for the model which is trained on all subdatasets except *i* and *j*. The data cube was then transformed to the normalized relative change:

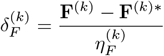

where **F**^(*k*)*^ is the performance of the control model on the subdataset *k* and 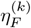 is again a normalization factor such that *δ*_*F*_ ∈ [−1, 1]. To quantify the similarity of our subdatasets we defined a redundancy metric which we call one-way redundancy *ρ*^1^:

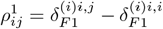

where F1 is taken from a model with subdatasets *i, j* removed from training and evaluated on subdataset (*i*), corresponding to the extra decrease in performance when omitting the second (*j*) on top of the first (*i*) subdataset. Since we want a symmetric redundancy metric in order to use it as a dissimilarity/distance metric, we then define the two-way redundancy *ρ*^2^ as the average of *ρ*^1^ across the diagonal:

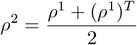

Both *ρ*^1^ and *ρ*^2^ ∈ [−1, 1] should be interpreted such that negative values correspond to synergism (positive performance cross-effect), while 0 corresponds to redundancy (neutral performance cross-effect) and positive values corresponding to antagonism (negative performance cross-effect).

## 3 Results

### 3.1 Experiment 1: Backbone size comparison

To provide a reasonable baseline for the performance of `flatbug` and a quantifiable justification when choosing a particular YOLOv8 backbone size with `flatbug`, we conducted a standard train-validation experiment with the four available backbone sizes: **<u>L</u>**arge, **<u>M</u>**edium, **<u>S</u>**mall, and **<u>N</u>**ano. All models in this experiment were trained for 300 epochs using our training dataset consisting of 23 subdatasets and 5153 images and evaluated on our evaluation dataset with the same subdatasets and 978 images.

Here, `flatbug` achieves cutting-edge performance of 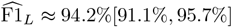 for the large `flatbug` variant and 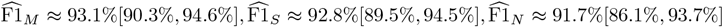 for the medium, small and nano `flatbug` variants respectively (numbers in square brackets are 95% confidence intervals). These metrics are concordant with the traditional machine-learning precision-recall curve and AP50%/AP50-95% metrics (Appendix L). As expected, smaller variants performed slightly worse on average, with more substantial decreases in worst-case performance of particularly the smallest of the variants; 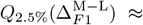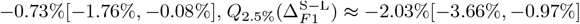 and 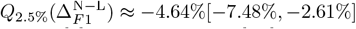 (Fig. 3). Upon close inspection (Fig. 3A), we found that differences between model sizes were primarily driven by a few outlier datasets, particularly ATO, BIS, and AMA in precision and AMI and BDA in recall (Fig. S6).

**Figure 3:**
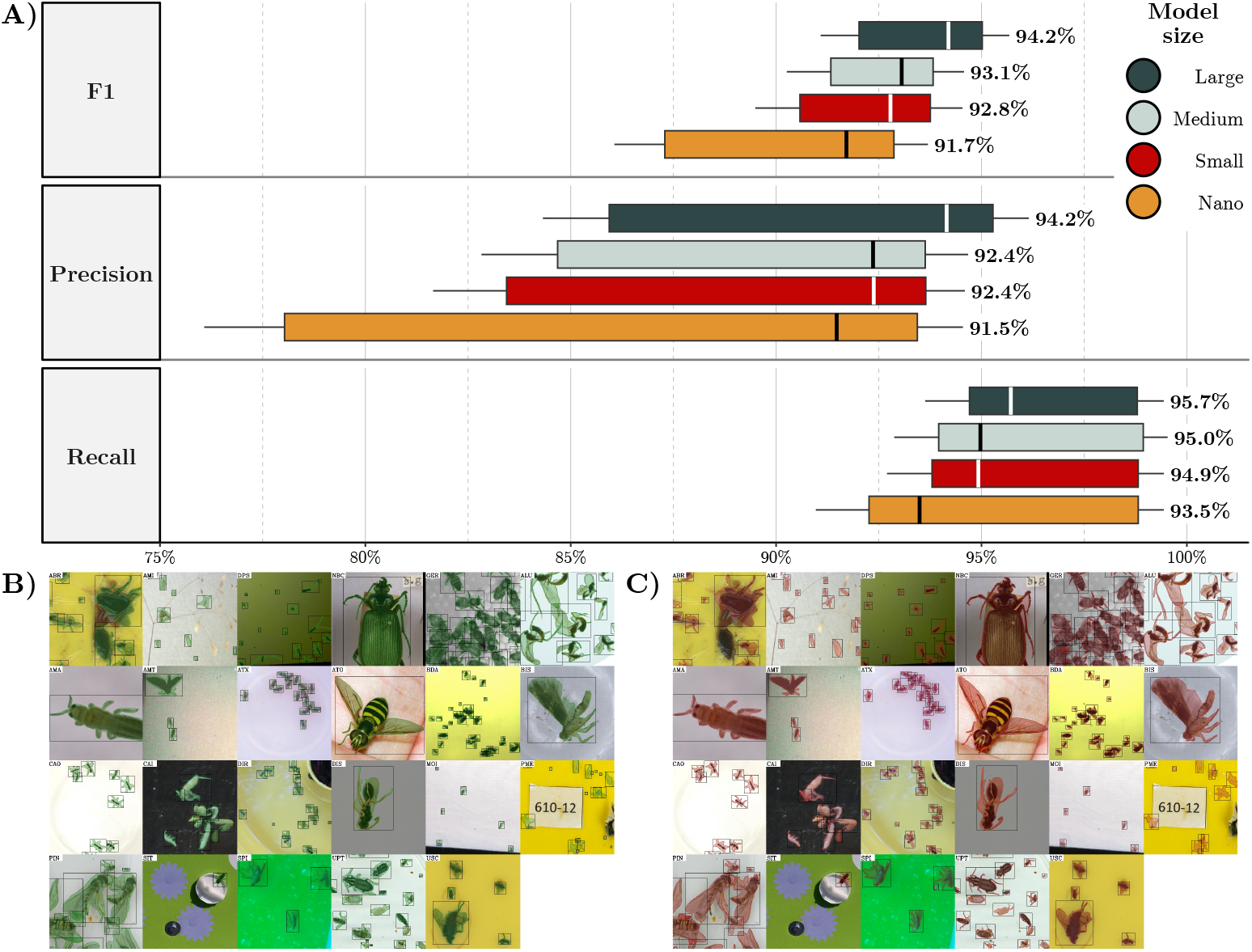
Validation performance of the deployment `flatbug` models at the four available model sizes (**A**) with true label (**B**) and predicted instance segmentation polygons and bounding boxes (**C**) on a hand-picked ROI (in the validation split) from each subdataset in the `flatbug` dataset. Box plots are based on a subdataset stratified bootstrap of model instance predictions (TP/FP/FN) on the validation split and show the 2.5%, 25%, 50% (median; also written next to each box plot), 75% and 97.5% bootstrap quantiles.

From a qualitative comparison of the true label instance polygons (Fig. 3B) the inferred instance segmentation polygons produced by the large (L) `flatbug` variant (Fig. 3C) also reveal high-fidelity contours and a low rate of false positives and negatives. The differences between the predicted and labeled detections can broadly be grouped into three categories; (1) the labeled instance is smaller than model size detection-threshold [AMI=1; DPS=1; BDA=1; DIR=1; PME=5], (2) incorrect separation of instances [CAI=1; PIN=3] and (3) instance not detected or non-instance incorrectly detected [ABR=1; AMI=6; SPI=1] (Fig. 3B-C). These results show that `flatbug` is a strong multiple-object detection and segmentation model for a broad spectrum of terrestrial arthropod images and instance sizes, and that particularly the three largest versions (L/M/S) are robust across the subdataset domains.

### 3.2 Experiment 2: Leave-one-out cross-validation

To assess the potential generalizability of `flatbug`, we investigated the out-of-box performance penalty in a leave-one-out cross-validation experiment. As explained in Section 2.5.2, leave-one-out models were trained on all subdatasets except one, for 50 epochs, subsequently the models were then evaluated only on the particular subdataset which was excluded (left-out) giving the leave-one-out performance. The leave-one-out models were then compared to a control model, which was trained on all subdatasets for 50 epochs. Then, each of the leave-one-out models was trained (fine tuned) for a further 10 epochs, only on the particular dataset which was excluded during the initial training. The control model was again trained (fine tuned) on all subdatasets, and compared with each fine-tuned leave-one-out model based on performance for only the relevant subdataset. Crucially, for both the leave-one-out and fine-tuned models, our comparisons include only the performance on the relevant subdataset for both the treatment and control model, not their overall performance on the whole dataset.

We identified a small, but significant, decrease in F1 performance of *δ*_F1_ ≈ −7.1%[−12.1%, −0.4%] when leaving individual subdatasets out. This was driven by a larger significant decrease in recall *δ*_R_ ≈ −10.3%[−13.9%, −6.6%], while precision remained unchanged *δ*_P_ ≈ 4.5%[−4.8%, 13.7%] (see Section 2.4 for details on *δ*). Although precision was not significantly affected in the leave-one-out scenario on average, there was significant variation among subdatasets, with a strong negative impact on subdatasets BIS 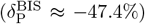 and SIT 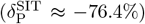, while conversely AMT 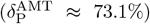 and CAO 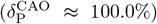 reached higher precision when excluded from training. We also found that fine-tuning these models for a modest number of epochs (10) on only the previously left-out dataset consistently removed the performance penalty *δ*_F1_ ≈ 0.9%[−1.3%, 6.7%], *δ*_R_ ≈ 0.0%[−2.4%, 4.5%], while marginally improving precision *δ*_P_ ≈ 5.1%[1.7%, 9.9%] (Fig. 4).

**Figure 4:**
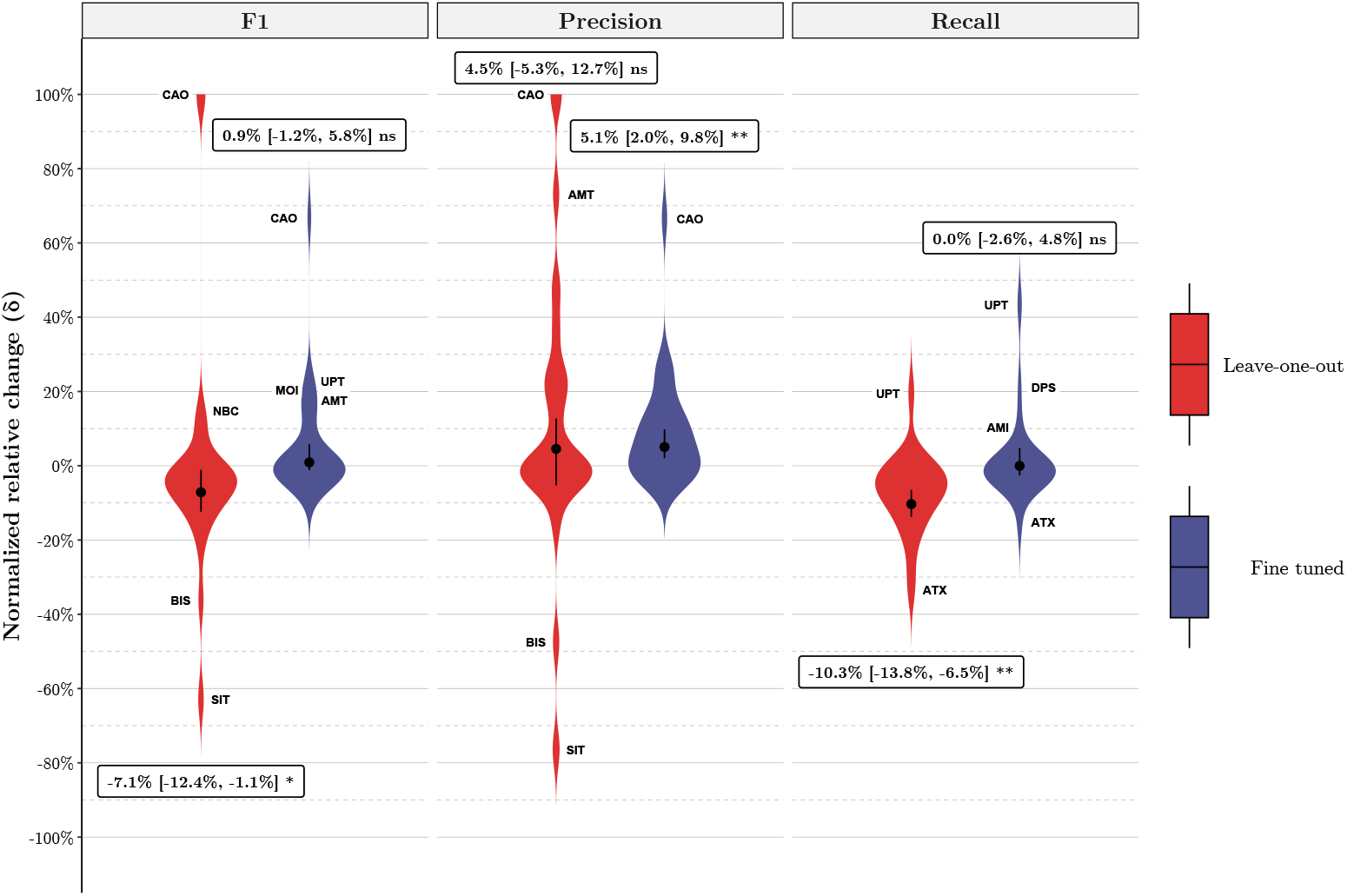
Density estimates (violins) and bias-corrected and accelerated (BCa)(DiCiccio & Efron, 1996) confidence intervals with empirical p-values(North et al., 2002) based on a weighted bootstrap for the normalized relative change, *δ*, (the proportion of the potential performance degradation/gain achieved under the treatment; see Section 2.4 for details on *δ*) between a leave-one-out and a control `flatbug` model (M) (leave-one-out) and a fine-tuned and a second control `flatbug` model (fine-tuned). Confidence interval and density for *δ*are weighted proportionally to the logarithm of the number of instances (true or false) in each subdataset as a compromise between micro- and macro-averaging. Three-letter annotations (e.g. CAO) are ‘outlier’ subdatasets for each sub-experiment. By definition, the normalized relative change, *δ*, of the control models is 0 in all cases, meaning that confidence intervals that overlap with zero, represent scenarios in which the performance of the alternate models are not significantly different from the control models.

This demonstrates that `flatbug` is potentially able to generalize to novel subdomains across most terrestrial arthropod images, even without any retraining, and that even a small amount of retraining—only including images from the novel domain—will lead to high performance, similar to that expected from a domain-specific model.

### 3.3 Experiment 3: Leave-two-out cross-validation

To explore our coverage and robustness across the potential data domain, we defined a pairwise subdataset redundancy metric—two-way redundancy, *ρ*^2^, (see Section 2.5.3 for details on *ρ*^2^)—based on the leave-two-out cross-validation experiment using 16 out of the 23 subdatasets in the complete `flatbug` dataset. We found that most (11 out of 16) of the investigated subdatasets hierarchically cluster in a single central synergistic cluster (measured by F1) interactions, while three subdatasets (ATO, DPS, CAI) are not positively affected by the other investigated subdatasets and the remaining two subdatasets (BIS, PIN) appear to form an outlier cluster with mutual benefit (Fig. 5). On the other hand, we found that the pair AMI-AMT is strongly synergistic with the lowest two-way redundancy, while being firmly placed in the major cluster, which should be expected as these two subdatasets originate from the same camera trap and imaging system (although from different iterations).

**Figure 5:**
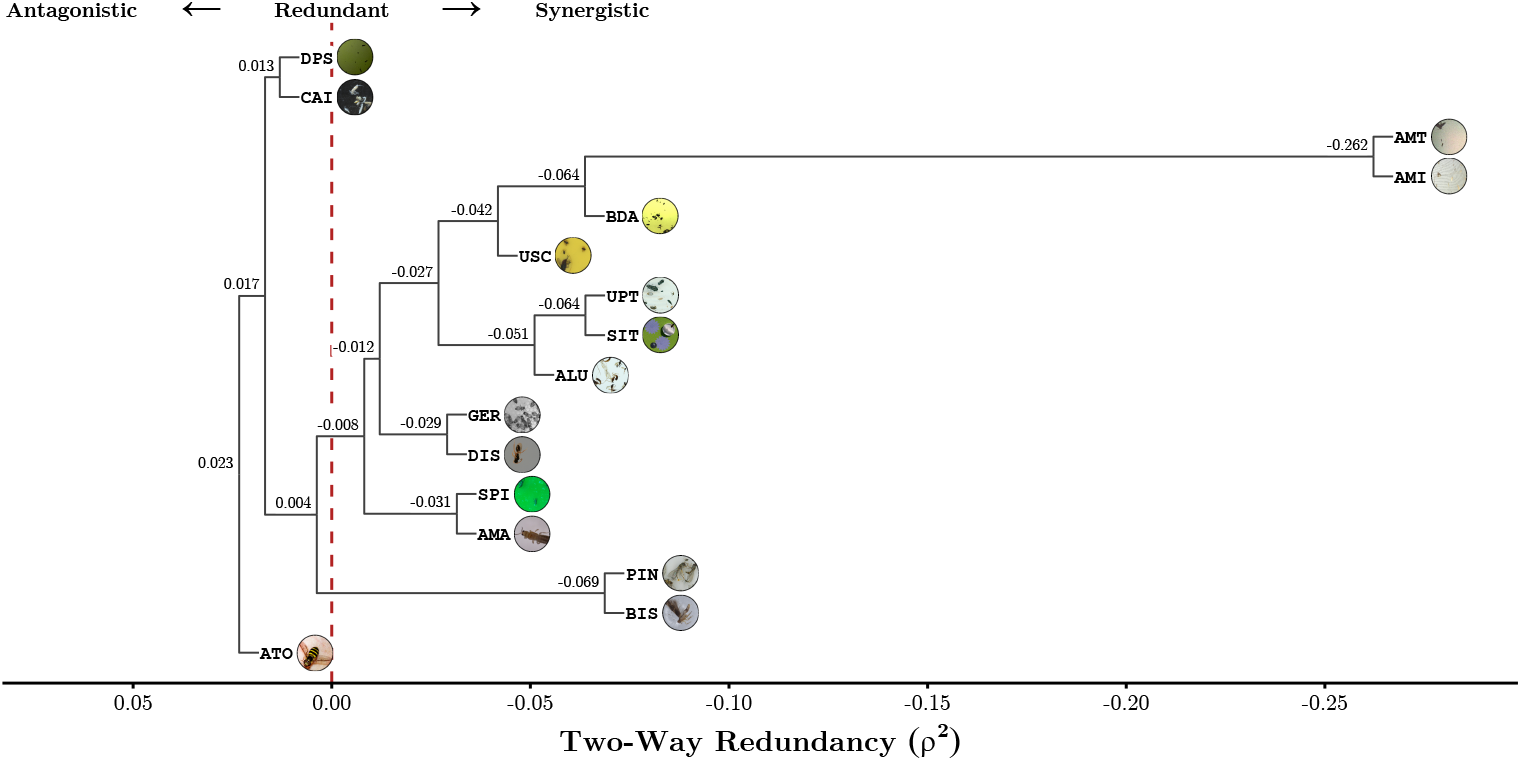
Hierarchical clustering of the pairwise two-way redundancy, *ρ*^2^, between 16 out of the 23 subdatasets based on F1 performance on the validation dataset. Negative values of *ρ*^2^ can be interpreted as synergism with a positive performance interaction between subdatasets, zero corresponds to perfectly redundant subdatasets with no performance interaction, while positive values correspond to antagonistic subdatasets with a negative performance interaction (see Section 2.5.3 for further details on *ρ*^2^). The `flatbug` performance on all subdatasets which are joined by an internal node below zero should be considered ‘resilient’, meaning that the model should be able to perform relatively well on other similar datasets.

We corroborated the overall pattern by an ordination of the pairwise redundancy matrix, this ordination has a similar, but not identical structure, to the hierarchical clustering tree, where subdatasets BIS, CAI and DPS qualitatively appear as outliers. By constructing a minimum spanning tree on the ordinated subdatasets, we also noted that most (10 out of 15) edges in the minimum spanning tree have negative two-way redundancies, with most (3/5) of the positive edges connect to ATO, DPS and CAI (Fig. S4), further validating the existence of a major central synergistic cluster, and the identity of the outlying subdatasets.

These patterns, particularly the existence of a major central synergistic cluster, show that the `flatbug` dataset (Geissmann & Svenning, 2025) has high coverage across its central domain (top-down images of terrestrial insects with low-to semi-complex backgrounds), while some of the peripheral domains (such as citizen science images) would benefit from the addition of similar subdatasets.

## 4 Discussion

We have presented `flatbug`, a powerful, general, scale- and size-agnostic model for precise detection and segmentation of individual arthropods from image-based approaches in entomology. It provides a simple and flexible out-of-the-box method for precisely extracting individual organisms in a very broad range of images.

As image-based approaches in entomology continue to gain traction (Høye et al., 2021; Kitzes et al., 2021; Schneider et al., 2023; van Klink et al., 2022), `flatbug` is solving the critical challenge of accurate and efficient arthropod detection. In combination with the `flatbug` accompanying dataset (Geissmann & Svenning, 2025) and robust performance evaluation, we pave the way for developments in individual detection within the automated arthropod monitoring community.

A major methodological milestone in computer vision for entomology is moving detection from specialized, bounding-box-based approaches to general, segmentation-based approaches. Previous methods for individual detection within ecological monitoring, particularly within terrestrial arthropod or entomological monitoring, and research have had limited taxonomic scope and little or no ability to generalize, whether to different taxonomic or image domains (Gal, Saragosti, & Kronauer, 2020; Geissmann, 2022; Hong et al., 2021; Kimura & Sota, 2023; Kloster et al., 2023; Sys et al., 2022; Z. Wu et al., 2023). Although some studies have attempted to segment arthropods in images (Geissmann, 2022), arthropod detection studies rarely attempt to demonstrate generality (Gal, Saragosti, & Kronauer, 2020; Hong et al., 2021; Sys et al., 2022), and tests of generalization, such as the leave-one-out experiment presented here, are extremely rare (Mazen, 2023; Willi et al., 2019).

We believe this discrepancy is caused by a lack of standardization between datasets and a lack of FAIR approaches to data management. Furthermore, significantly reduced annotation effort for segmentation masks compared to bounding boxes has meant that the potential benefits of segmentation is underexplored and likely understated.

We argue that segmentation offers four major benefits: (1) the background of detected individuals can be removed, reducing potential bias in downstream classification, (2) inference deduplication with non-max suppression of segmentation polygon IoU offers a more accurate estimate for instance overlap compared with bounding box IoU, (3) individual size estimation is much more accurate using segmentation areas than bounding box areas, especially for long-legged arthropods and (4) it opens up the possibility for semantic segmentation of individual body parts as in Mráz et al. (2024).

`flatbug` contributes to overcoming these methodological problems in multiple ways, where the three main contributions to improving the localization of arthropods in images can be summarized as instance representation, input size independence, and dataset diversity and structure.

The first key feature of `flatbug`, is the polygon-based instance representation through our strict adherence to a segmentation-first approach for the entire inference pipeline. Whereas YOLOv8 utilizes bounding boxes for deduplication (Jocher et al., 2023), `flatbug` uses the segmentation contour polygons for deduplication. To overcome and essentially trivialize the significant additional overhead associated with computing the intersection of arbitrary simple 2D polygons (segmentation contours), compared to the intersection of axis-aligned rectangles (bounding boxes), `flatbug` introduces a novel two-stage NMS (non-max suppression) algorithm.

The second key feature of `flatbug` is our solution to the scale-dependence of fixed-size inference detection models like YOLOv8 (Jocher et al., 2023), and even the hyper-inference framework SAHI (Akyon et al., 2022), that is often used to alleviate this issue. To solve this problem our hyper-inference algorithm employs a ‘tile-pyramid’ approach, tiling the image at geometrically spaced scales—from the native scale to the full image—using fixed-size tiles with specified overlap (Fig. 2). This differs from SAHI’s single-scale tiling by allowing detection at multiple scales, improving the identification of objects of varying sizes. Our method enables scale-agnostic detection on very large images, accurately identifying all instance sizes—from the smallest visible arthropods to the largest—including intermediate sizes, rather than being limited to only the extremes.

The third key feature of `flatbug`, is the uniquely large and diverse, in terms of taxonomic coverage and image types, segmentation training dataset (Geissmann & Svenning, 2025) that forms the foundation of the `flatbug` model(s). The unique structure, size, and diversity of our dataset allows us to investigate critically needed generalized performance in an unprecedented manner, enabling robust claims about the out-of-box and out-of-domain performance. Previous methods have not been able to effectively quantify performance on novel data captured using different imaging systems and/or with different taxa, due to more narrow test datasets, often simply consisting of a subset of their specialized dataset. We show that `flatbug` can be directly applied in new contexts (without retraining) by utilizing the explicit subdomain stratification of our dataset in a leave-one-out cross-validation experiment, with only a minor performance degradation of -7.1% [-12.1%, -0.4%] relative decrease in F1 (*δ*_F1_) on most datasets (Section 3.2).

The variation in performance degradation between subdatasets in the leave-one-out experiment (Fig. 4) suggests an underlying structure and similarity between subdatasets. To investigate this similarity, we expanded our analysis with a pairwise leave-two-out experiment to define a subdataset dissimilarity metric to map the domain space. In order to explicitly quantify the similarity structure, we design the novel dissimilarity metric ‘two-way redundancy’ (*ρ*^2^), which describes how the performance of `flatbug` on one dataset changes, when another dataset is removed along with it. Of the 16 subdatasets we investigated in this analysis, we found that most datasets form a major synergistic cluster, with the subdatasets AMI and AMT being particularly synergistic, as is expected since these subdatasets originate from the same imaging system (Fig. 5). On the other hand, the subdatasets ATO, DPS and CAI appeared to be redundant to antagonistic with the remaining subdatasets, suggesting a considerable domain shift. This is likely explained by ATO being the only subdataset consisting of diverse images of insects on natural backgrounds (citizen science images), while CAI is unique in containing images of white insects (Collembola) on a black background (soil). It was somewhat surprising that DPS emerged as an outlier, given that its images are captured by a light-attracting moth camera trap, from which we expected it to cluster with AMI and AMT. We speculate that slightly lower-quality images on a yellowish screen (instead of white), combined with lower labelling consistency, might explain this discrepancy.

The observed generalizability and its boundaries offer a direct and simple path to improving `flatbug`, and set the stage for future research within machine learning in automated arthropod monitoring. Through community inclusion and efforts, additional datasets and annotations will likely be able to further improve `flatbug`, especially as more of the vast array of citizen science images from public platforms can be integrated. While `flatbug` is built on YOLOv8, much of our work and code is model-agnostic and can be adapted to emerging architectures. For example, YOLOv11—the successor to YOLOv8—has already been released, a trend we expect to continue. We have also demonstrated that although hyper-inference frameworks such as SAHI already exist, there is considerable scope for improvements in this area. For the method we have presented, this is particularly evident in the `flatbug` pyramid-tiling algorithm produces a very high degree of tile redundancy (Fig. 2), which could be reduced by further research, possibly leading to a significant decrease in the computational cost of inference.

`flatbug` is an open-source Python package (https://github.com/darsa-group/flat-bug/), which we believe offers cutting-edge performance and user-friendliness in its current state, due to its ease of installation, use and integration through our Python and CLI API and online demo (https://colab.research.google.com/github/darsa-group/flat-bug/blob/master/docs/flat-bug.ipynb). Crucially, unlike previous approaches `flatbug` is scale- and size-agnostic, has a broad taxonomic domains and can tackle a wide diversity of image types, although we also outline several key paths for improvement. As our experiments show, through multiple novel technological developments, `flatbug` as a tool opens up several new research avenues in entomology, conservation, monitoring, agriculture, and beyond.

## 5 Acknowledgements

We are thankful to a number of contributors who collected and provided previously unpublished images (and annotations) for specific sub-datasets (denoted by their three-letter code), including Nathan Pinoy; Paul Abram(ABR); Hans Jørgen Skydt Andersen and Jeppe Fogh Rasmussen(BDA); Mariana Abarca(MOI); and Tim Gernat and Gene Robinson(GER). We would also like to acknowledge testers of flatbug for their valuable feedback and help in improving the initial versions, these include: Graham Smith, Sara Nawoya, Tom August and his group, and David Rolnick and his group. The work was supported by the Global Innovation Network Program grant no. 2084-00048 (Danish Ministry of Higher Education and Science). Part of the computation done for this project was performed on the UCloud interactive HPC system, which is managed by the eScience Center at the University of Southern Denmark. The work was supported by the European Union’s Horizon Europe Research and Innovation programme, under Grant Agreement No. 101060639 (MAMBO) and is partly based upon work from COST Action InsectAI CA22129, supported by COST (European Cooperation in Science and Technology). Work in the Plant Insect Ecology and Evolution Lab (J.C., D.C., N.C.) was supported by the Natural Sciences and Engineering Research Council of Canada (NSERC), ALLRP 570736-2021, and the AgriScience Program under Agriculture and Agri-Food Canada’s Sustainable Canadian Agricultural Partnership as part of Organic Science Cluster 4. Q.G. was supported by the Novo Nordisk Foundation Start Package grant (NNF22OC0077040) and the 2022-2023 BIODIVERSA+ joint call for research proposals, with the funding organisation Innovation Fund Denmark (grant no. 2128-00003).

## 7 Author Contributions

A. Svenning, Q. Geissmann and T. T. Høye conceived the ideas. A. Svenning, G. Mougeot and Q. Geissmann contributed to the implementations. A. Svenning, G. Mougeot, Q. Geissmann and T. T. Høye led the writing. A. Svenning, D. Chevalier, G. Mougeot, J. Alison, J. Carrillo, K. Bjerge, Q. Geissmann, S. -Q. Ong and T. T. Høye contributed significantly to review and editing of the drafts. A. Svenning and G. Mougeot analysed the results. A. Svenning, N. C. Molina, N. Pinoy, Q. Geissmann and S. -Q. Ong contributed to training and evaluation data annotation. D. Chevalier, J. Alison, J. Carrillo, N. C. Molina, Q. Geissmann and S. -Q. Ong collected the data. J. Carrillo, Q. Geissmann and T. T. Høye were instrumental in project management and supervision. All authors contributed critically to the drafts and gave final approval for publication.

## 8 Data Availability

Data available on Zenodo https://www.doi.org/10.5281/zenodo.14761447 (Geissmann & Svenning, 2025). Code is available on GitHub https://github.com/darsa-group/flat-bug/.

Appendix

### J Dataset content description

For ease-of-access we provide DOIs and references for both our full `flatbug` dataset, as well as each component subdataset in Table S1. Since many of the subdatasets have been used in association with a previous study, as well as having been published as a standalone dataset, we provide references to both the research article, as well as the dataset publication, and also provide a new standardized DOI for each subdataset. The standardized DOIs serve three purposes; (1) they allow citation of single subdatasets which have not previously been published, (2) they specifically cite the version of the subdatasets we used and (3) they are all hosted along with the data on Zenodo, providing a simple unified access GUI and API for all subdatasets (as well as the full `flatbug` dataset).

**Table S1:** Subdataset data, reference and source.

|  | DOI |  |  |
| --- | --- | --- | --- |
|  | Data | Article | Data Source |
| ABR | [10.5281/zenodo.13992282](https://doi.org/10.5281/zenodo.13992282) | - | - |
| ALU | [10.5281/zenodo.13919486](https://doi.org/10.5281/zenodo.13919486) | [10.1111/2041-210X.13769](https://doi.org/10.1111/2041-210X.13769)[44] | [10.5683/SP2/LMRVFN](https://doi.org/10.5683/SP2/LMRVFN)[43] |
| AMA | [10.5281/zenodo.13919602](https://doi.org/10.5281/zenodo.13919602) | [10.1016/j.compag.2022.107462](https://doi.org/10.1016/j.compag.2022.107462)[4] | [10.26180/21547650.v3](https://doi.org/10.26180/21547650.v3)[3] |
| AMI | [10.5281/zenodo.13991167](https://doi.org/10.5281/zenodo.13991167) | - | - |
| AMT | [10.5281/zenodo.13991268](https://doi.org/10.5281/zenodo.13991268) | [10.1111/icad.12662](https://doi.org/10.1111/icad.12662)[36] | [10.17192/fdr/194](https://doi.org/10.17192/fdr/194)[35] |
| ATX | [10.5281/zenodo.13919641](https://doi.org/10.5281/zenodo.13919641) | [10.7554/eLife.58145](https://doi.org/10.7554/eLife.58145)[13] | [10.5281/zenodo.3740546](https://doi.org/10.5281/zenodo.3740546)[12] |
| ATO | [10.5281/zenodo.13919717](https://doi.org/10.5281/zenodo.13919717) | [10.1186/s44147-023-00284-8](https://doi.org/10.1186/s44147-023-00284-8)[34] | [10.34740/kaggle/dsv/1240192](https://doi.org/10.34740/kaggle/dsv/1240192) |
| BDA | [10.5281/zenodo.13992340](https://doi.org/10.5281/zenodo.13992340) | - | - |
| BIS | [10.5281/zenodo.13920322](https://doi.org/10.5281/zenodo.13920322) | [10.48550/arXiv.2307.10455](https://doi.org/10.48550/arXiv.2307.10455)[17] | [10.5281/zenodo.8015206](https://doi.org/10.5281/zenodo.8015206)[18] |
| CAO | [10.5281/zenodo.13920372](https://doi.org/10.5281/zenodo.13920372) | [10.1093/gigascience/giac096](https://doi.org/10.1093/gigascience/giac096)[61] | [10.17632/9ws98g4npw.3](https://doi.org/10.17632/9ws98g4npw.3)[8] |
| CAI | [10.5281/zenodo.13920408](https://doi.org/10.5281/zenodo.13920408) | [10.1111/2041-210X.14001](https://doi.org/10.1111/2041-210X.14001)[50] | - |
| DPS | [10.5281/zenodo.13991200](https://doi.org/10.5281/zenodo.13991200) | - | [10.5281/zenodo.10853097](https://doi.org/10.5281/zenodo.10853097)[22] |
| DIR | [10.5281/zenodo.13992477](https://doi.org/10.5281/zenodo.13992477) | [10.3390/jimaging4110129](https://doi.org/10.3390/jimaging4110129)[28] | - |
| DIS | [10.5281/zenodo.13929922](https://doi.org/10.5281/zenodo.13929922) | [10.1111/1755-0998.13567](https://doi.org/10.1111/1755-0998.13567)[64] | [10.17605/OSF.IO/EN594](https://doi.org/10.17605/OSF.IO/EN594)[63] |
| GER | [10.5281/zenodo.13991237](https://doi.org/10.5281/zenodo.13991237) | - | - |
| MOI | [10.5281/zenodo.13992498](https://doi.org/10.5281/zenodo.13992498) | - | - |
| NBC | [10.5281/zenodo.13991316](https://doi.org/10.5281/zenodo.13991316) | - | [10.5519/0018095](https://doi.org/10.5519/0018095)[26] |
| PME | [10.5281/zenodo.13930055](https://doi.org/10.5281/zenodo.13930055) | - | - |
| PIN | [10.5281/zenodo.13992379](https://doi.org/10.5281/zenodo.13992379) | - | - |
| SIT | [10.5281/zenodo.13920464](https://doi.org/10.5281/zenodo.13920464) | - | [10.5281/zenodo.7725940](https://doi.org/10.5281/zenodo.7725940)[48] |
| SPI | [10.5281/zenodo.11366139](https://doi.org/10.5281/zenodo.11366139) | - | [10.5281/zenodo.6382496](https://doi.org/10.5281/zenodo.6382496)[14] |
| UPT | [10.5281/zenodo.13992496](https://doi.org/10.5281/zenodo.13992496) | - | - |
| USC | [10.5281/zenodo.13992488](https://doi.org/10.5281/zenodo.13992488) | - | - |

### K Deterministic oversampling

We calculate initial oversampling weights for all images, *ω*^*^:

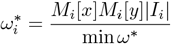

where *M*_*i*_[*q*] is the size of the image *i* in dimension *q* and |*I*_*i*_ | is the number of labeled instances in image *i*, such that the minimum initial weight is 1: min(*ω*^*^) = 1. The initial weights are chosen such that larger images with a high density of instances are oversampled more than smaller images with fewer instances. Then we calculate the final oversampling weight, *ω* (denoting how many times we see an image in a single epoch), given an inflation factor, *λ* (default=2) (we target *λ* times the number of images, as the total number of training samples per epoch):

#### Algorithm S2

Deterministic oversampling factor for re-balancing strata.

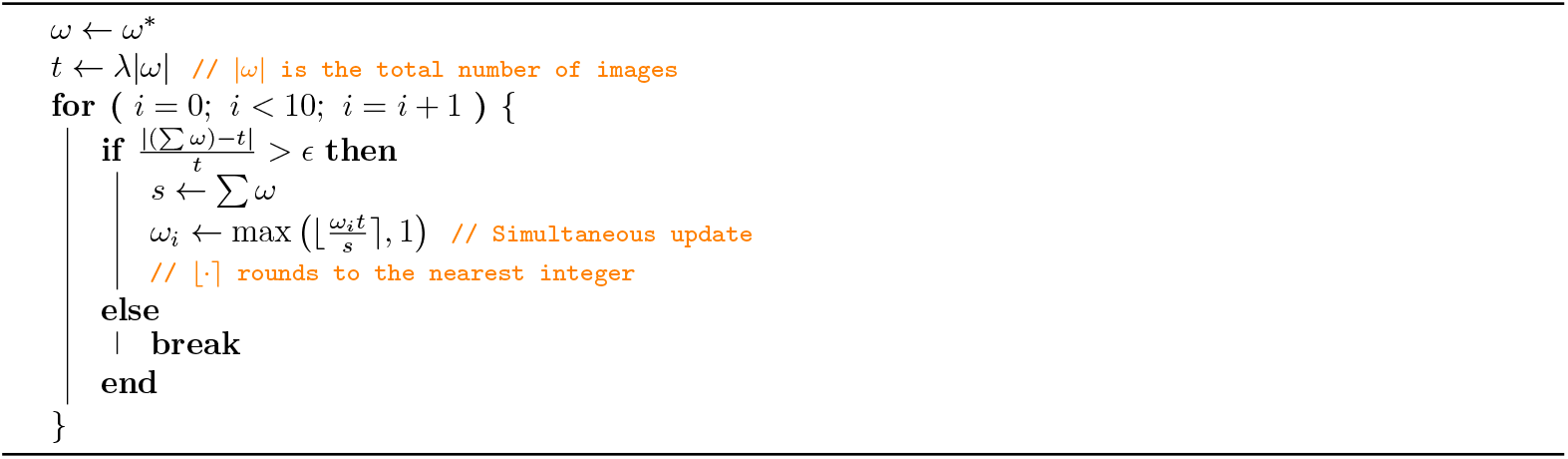

The purpose of the design in Algorithm S2 is to compute a list of positive non-zero integers, *ω*, that are approximately proportional to a given list of positive non-integers, *ω*^*^. In other words, each entry of *ω* is a scaled version of the corresponding entry of *ω*^*^ by roughly the same factor, subject to the constraint that every entry must be an integer greater than or equal to 1. At the same time, the total sum ∑*ω* is kept close to a target *t*, where *t* is defined as a multiple, *λ*, of the length of the list: *t* = *λ* |*ω* |. The number of iterations (10) is chosen arbitrarily, empirical observations verify that the algorithm converges very fast, typically achieving a sum very close to the target after only a few rounds. These integers are then used to create a random oversampled permutation of the images in the training data during an epoch, while guaranteeing that we see all images at the exact same frequency every single epoch.

### L Experiment 1: Precision-Recall Curve and Average Precision

For those readers who are more accustomed to precision-recall curves and AP50%/AP50-95%, we have also computed these for the deployment models at the four YOLOv8 backbone sizes from experiment 1.

As we also showed in Section 3.1, across metrics, including both AP50% and AP50-95%, the larger models in general perform somewhat better: AP50%_*L*_ ≈ 0.937, AP50%_*M*_ ≈ 0.932, AP50%_*S*_ ≈ 0.933, AP50%_*N*_ ≈ 0.914 and; AP50 − 95%_*L*_ ≈ 0.872, AP50 − 95%_*M*_ ≈ 0.862, AP50 − 95%_*S*_ ≈ 0.86, AP50 − 95%_*N*_ ≈ 0.821, with particularly the smallest model (N), losing significant performance, and the largest model (L), gaining some performance over the intermediate models (M/S). These results are mirrored in the precision-recall curve, with the curves for each model generally following eachother from 0-50% recall, where again especially the smallest model (N), start to significantly decrease compared to the others, while the intermediate models (M/S), decrease slightly more than the largest model (L) (Fig. S3).

**Figure S3:**
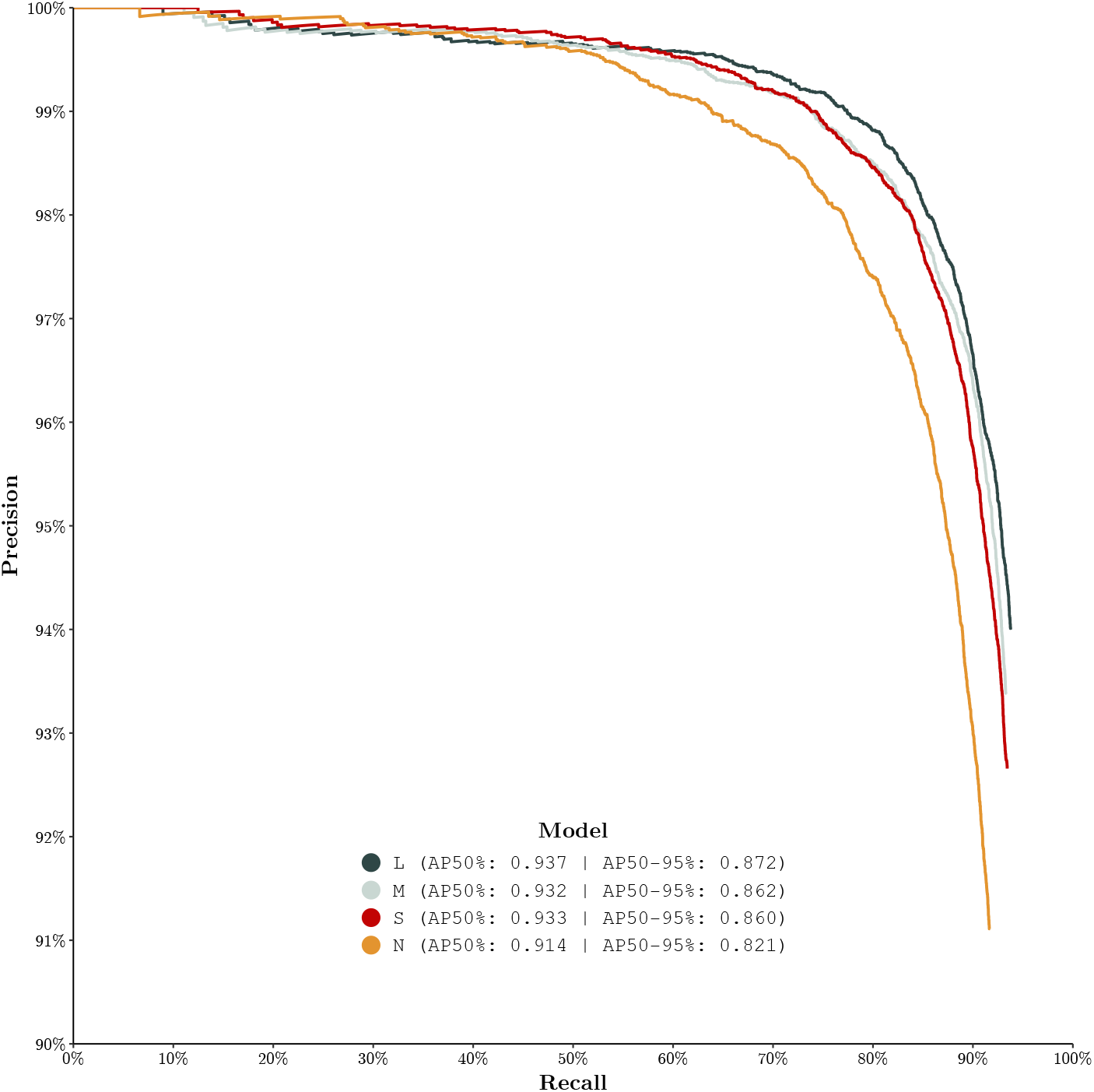
Precision-Recall curves for the four deployment models from experiment 1. Each model corresponds to one of the available YOLOv8 backbone sizes (**<u>L</u>**arge, **<u>M</u>**edium, **<u>S</u>**mall, and **<u>N</u>**ano). The legend is annotated with AP50% and AP50-95%, evaluated at 0.01 IoU threshold intervals.

### M Experiment 3: Leave-two-out cross-validation (addendum)

The two-way redundancy matrix, *ρ*^2^, can interpreted as a dense edge-weight matrix of an undirected graph with nodes for each of the 16 included in the leave-two-out experiment. Such a graph may not necessarily be projected into N dimensions, while maintaining that the euclidean distance matrix between the projected nodes is identical to the original edge-weight matrix, and as such any attempt to visualize this matrix in a figure such as Fig. 5, will come with assumptions, drawbacks and advantages. Therefore, we opted to add a secondary analogous, but methodologically different projection-based visualization to validate the patterns. While hierarchical clustering can accurately portray the relative ordering of relationships, it does not attempt to minimize loss of information about all edge-weights. Therefore, a logical counterpart to hierarchical clustering is a distance-based projection method such as PCoA, which from a given distance matrix, *d*, finds points in N dimensions, which have an euclidean distance matrix which approximates *d*. In our case we can easily transform *ρ*^2^ into a “distance”: *D*(*ρ*^2^) = (*ρ*^2^ +1)*/*2, and produce Fig. S4.

**Figure S4:**
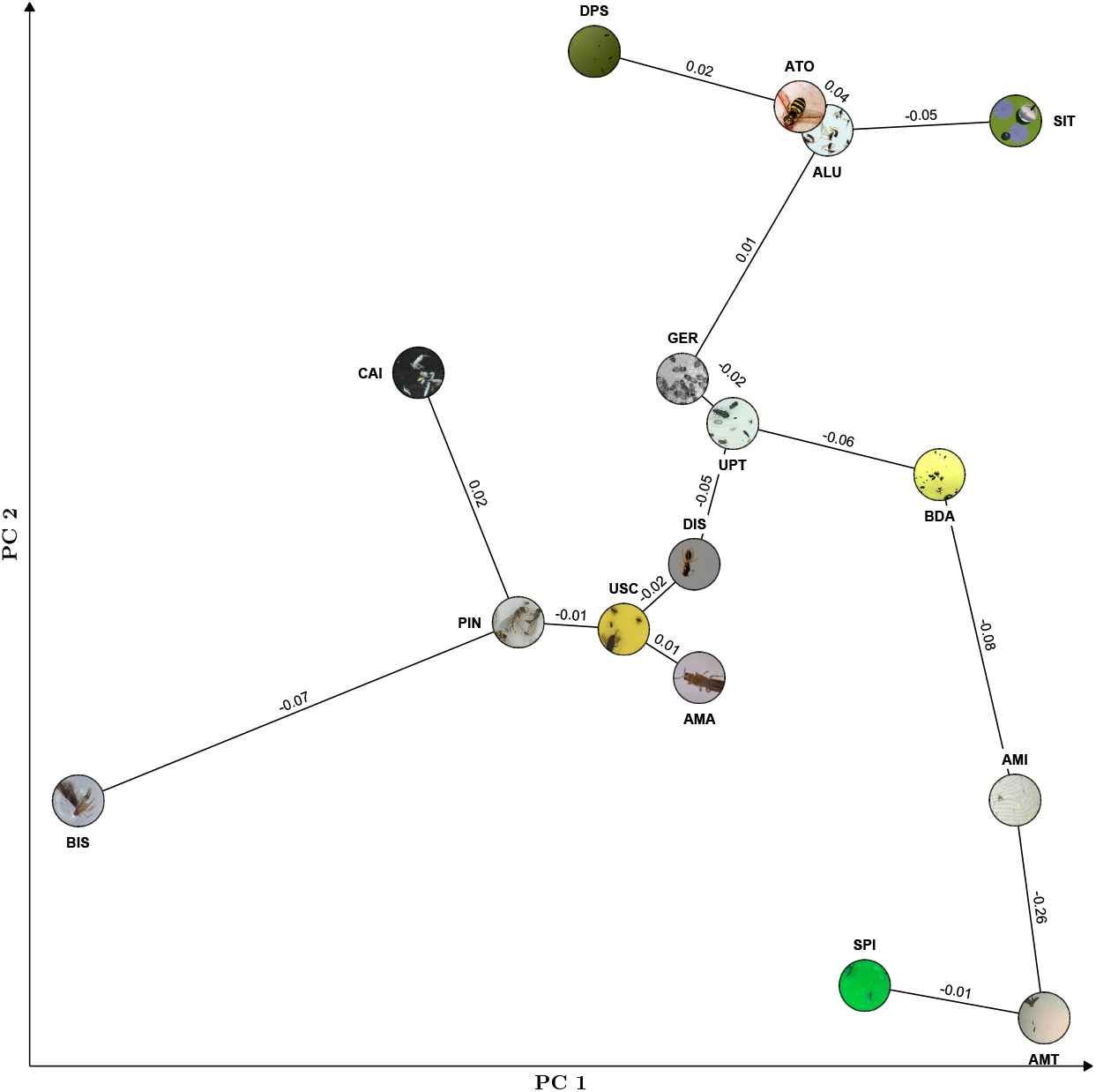
PCoA ordination of the pairwise two-way redundancy, *D*(*ρ*^2^) = (*ρ*^2^ +1)*/*2, matrix based on the leave-two-out cross-validation experiment using 16/23 of the `flatbug` subdatasets. The ordination is annotated with the euclidean minimum spanning tree in the 2D ordinated space, where the edges are labelled with the two-way redundancy (*ρ*^2^) between the connected subdatasets.

### N Statistical post-processing

For multiple reasons, we chose 32px (square-root contour-polygon area) as the cutoff, for the minimum object/instance size, considered during all stages, training, evaluation and inference. The reasons for this can be roughly grouped into three boxes: (1) practical and (2) epistemological. For practical reasons there are consistency and efficiency constraints to labelling, where as instances become smaller, the quality and consistency of labels decrease. On top of that, as instances become smaller, it becomes increasingly harder, even for the best expert labellers, to distinguish background from object, due to constraints on the information which can be contained in a (small) finite pixel-patch.

The exact cutoff, when it becomes impractical to continue labelling, and object and background become epistemologically identical, is beyond the scope of this paper, however we empirically observed that, for most of our datasets, the relative frequency of true positives decrease sharply around 32 pixels in square-root contour-polygon area (see Fig. S5).

**Figure S5:**
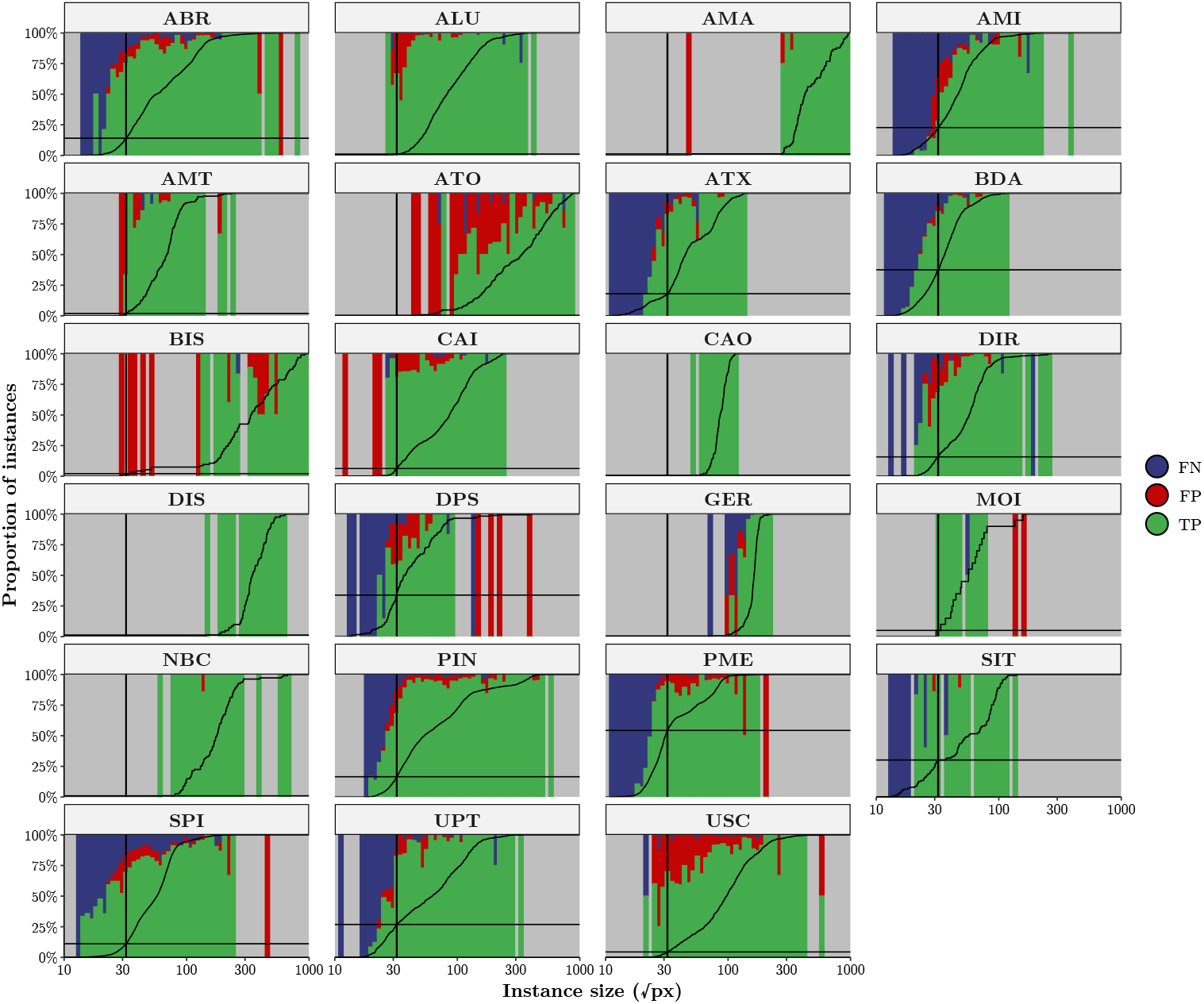
Frequency of true positives (TP), false positives (FP) & false negatives (FN) by instance size, stratified by subdataset. Instance size size is defined as the square root of the instance area in pixels (ground truth label are used for true positives and false negatives, and predictions for false positives). The vertical/horizontal black lines separates *“small”* instances—with a size less than 32 pixels—from *“large”* instances and shows which percentage of instances are *“small”*. The increasing black line shows the cumulative distribution of instances by size, that is its’ height indicates how many of the instances for a given dataset are at or below a given size.

### O Extra figures

**Figure S6:**
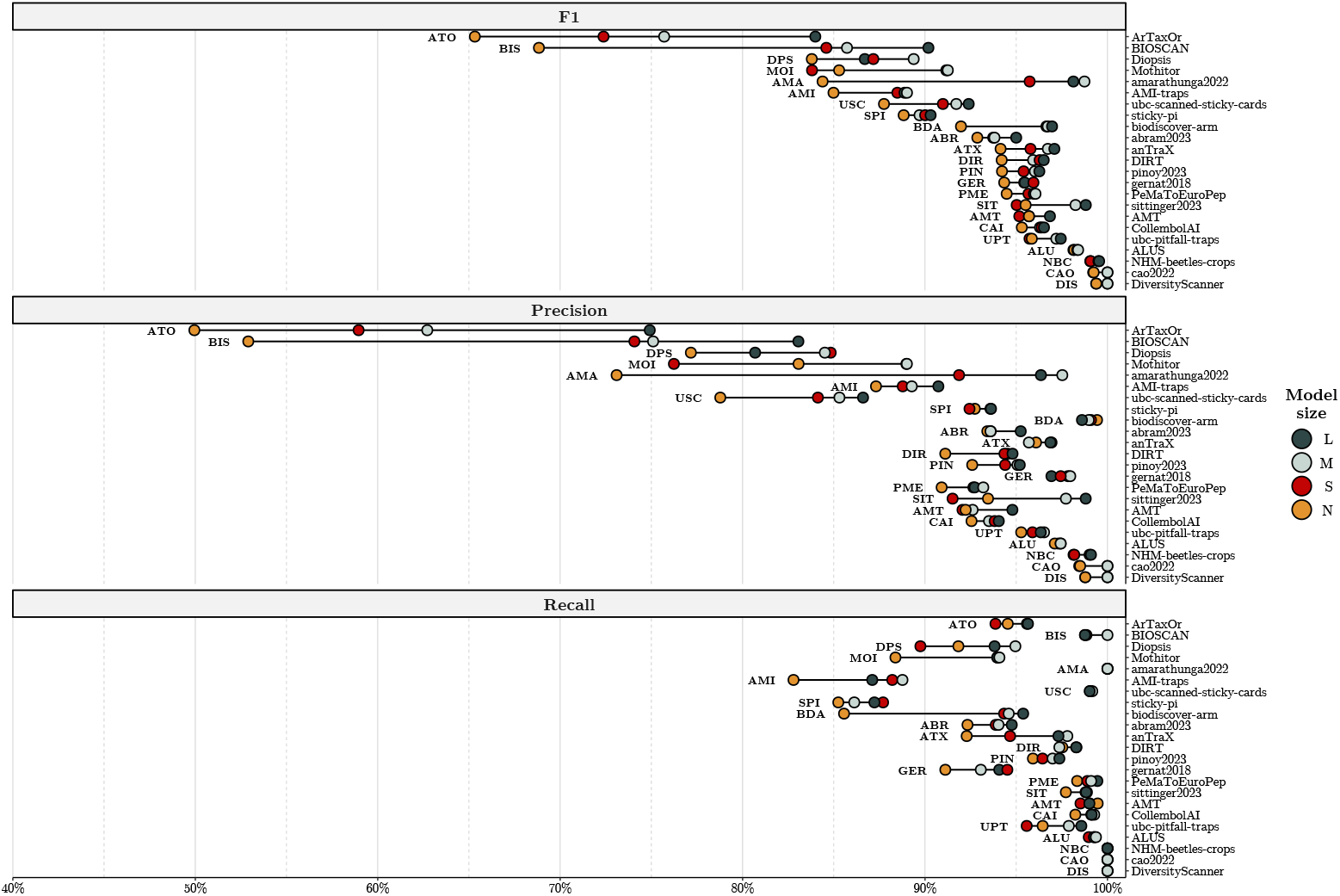
F1, recall and precision for our 4 deployment-ready `flatbug` models on each subdataset. Sub-datasets are ordered by minimum F1-score across model sizes.

**Figure S7:**
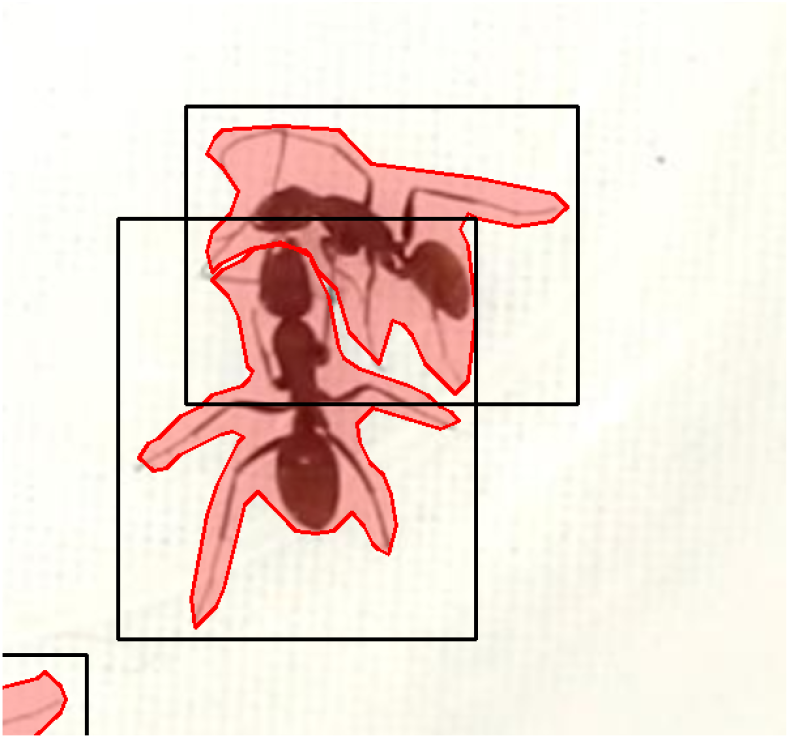
An example of segmentations being completely separate, while the bounding have significant overlab. This example clearly shows how segmentation polygons are strictly better at estimating object overlap (intersection).

